# Inquiry in Ridding the Democratic Republic of the Congo of sleeping sickness, a dream at our fingertips: comparing to the Epidemiology of human African trypanosomiasis in the Democratic Republic of the Congo 2002-2003

**DOI:** 10.1101/522771

**Authors:** Guyguy Kabundi Tshima

## Abstract

**Background:** In the Democratic Republic of the Congo, the international support was suddenly withdrawn after the massacre of students at the Lubumbashi University in May 1990. The interruption of the international aid from 1990 to 1991 would undoubtedly have a long-lasting negative effect on case load. So, the National Sleeping Sickness Control Programme—NSSCP (*Programme National de Lutte contre la Trypanosomiase Humaine Africaine*) (PNLTHA) remains vulnerable without international aid. Currently, the number of reported new cases decreased. These achievements prove that the elimination of this neglected tropical disease is possible when there is a strong commitment of public authorities accompanied by scientific research centers, civil society and the private sector. Without international aid, sleeping sickness remains a formidable disease difficult to cure because it is depending on continued financial support and drug availability.

**Objectives:** The objectives of this work were: 1. to profile the incidence of new cases of human African trypanosomiasis in the Democratic Republic of the Congo from 2002 to 2003, depending on the stage of disease progression (stages 1 and 2); 2. Compare the evolution of this profile from one household to another; 3. Compare the rate of confirmed parasitological diagnosis with positive CATT; 4. Calculate the discrepancy rate between CATT+ and parasitological diagnosis. All the above objectives are aiming to sustain the efforts made for the adoption of the 2018 Francophonie resolution on the ridding of human African trypanosomiasis that may renewed donor interest, including the government of the Democratic Republic of the Congo because the control of HAT is completely dependent on international aid.

**Methods:** Research is necessary on how to rationalize control activities so that control programs can adopt the most effective and efficient strategies. To assess it, we analyzed epidemiologic data collected by PNLTHA from 2002 to 2003.

**Results:** In all endemic areas, 1,970,101 people were tested in 2002 and 2,311,507 people in 2003. The national average coverage of the total population tested (TPT) represents 16.20% of the exposed population among which 13,853 new cases were detected in 2002 and 11, 481 new cases detected in 2003 with the national average coverage of the population tested that represents 19.10 %.

**Conclusion:** In short, we said that the number of people already infected is probably higher than the new cases reported in 2003. We are still far from the situation of 1958/60 when there was 1 new case declared by 10,000 people tested (i.e. 1,100 new cases out of 13,000,000 people screen). Therefore, the Congolese government must make long-term financial commitments to ensure the continuity of HAT control activities.

**Author summary:** For the past three decades, the frequency of sleeping sickness tends to become a large in the Democratic Republic of the Congo. This paper reviews the status of sleeping sickness in DRC between 2002 and 2003, with a focus on stage patterns. Epidemiological trends at the national and provincial level are presented. Today, this deadly fly-borne disease threatens more than 65 million people worldwide and most of the reported cases (more than 8 out of 10) are in the Democratic Republic of the Congo. Fortunately, after decades of hard work, we have never been so close to eradicating sleeping sickness in the Democratic Republic of the Congo. In 2009, the number of reported cases fell below 10,000, the first in half a century. In 2015, only 2,804 cases had been listed. The Democratic Republic of the Congo is determined to eradicate the disease by 2020, paving the way for its global eradication. Thanks to these decades of work. I submit this inquiry to recognize also the work that my last mentor coauthor of this search did, he hold his doctorate on this disease. This submission is an **appropriate way I found to honor and keep the memory of my supervisor who passed away!** In advance, many thanks for your best understanding of this particular circumstance. From my last mentor work, we have never been so close to the definitive elimination of sleeping sickness. The number of reported new cases decreased from 26,318 in 1998 to 11,481 in 2003 and later to 2,804 in 2015. These achievements prove that the elimination of neglected tropical diseases is possible when there is a strong commitment of public authorities accompanied by scientific research centers, civil society and the private sector.

## Introduction

Trypanosomiasis is an endemo-epidemic disease caused for Africa by trypanosoma gambiense and rhodesiense.

It is transmitted from a parasitized individual to a healthy individual by the bite of a specific vector insect, the tsetse flies or tsetse fly.

The infection has a fatal evolution for lack of a specific treatment, it begins with the invasion of the blood and the lymphatic ways by the parasite and continues after a variable delay by the attack of the nerve centers of the encephalon [1].

Currently, sleeping sickness tends to become a great endemic by its frequency, it remains a formidable disease difficult to cure.

Another important fact is the appearance in the cities of this disease, considered until now rural [2].

In the Democratic Republic of the Congo, the break-up of Belgian-Congolese cooperation in 1990 would undoubtedly have hindered the national program for the control of human African trypanosomiasis (PNLTHA).

### Research objectives

The objectives of this work were:

1. To profile the incidence of new cases of human African trypanosomiasis in the Democratic Republic of the Congo from 2002 to 2003, depending on the stage of disease progression (stages 1 and 2)
2. Compare the evolution of this profile from one household to another
3. Compare the rate of confirmed parasitological diagnosis with positive CATT
4. Calculate the discrepancy rate between CATT + and parasitological diagnosis

### Research hypothesis

The characterization of this evolutionary profile will make it possible to evaluate the effectiveness of the national program against trypanosomiasis (PNLTHA). In the case where this action is effective, the rate of new cases (NC) in stage 2 must be able to decrease significantly.

## Material and methods

### Material

We used the database established by the National Program for the Control of Human African Trypanosomiasis in the DRC (PNLTHA) of 2002 and 2003.

### Methods

This work consisted in the counting of the reports of the national program against trypanosomiasis (PNLTHA) of 2002-2003 concerning the provinces of the Democratic Republic of the Congo with an accent for the provinces mentioned like centers of the development of *trypanosomiasis* notably Equateur, Kasaï, Bandundu, Bas -Congo, Maniema, Katanga while noting the following:

- The location (district or province) having a high incidence
- The years: 2002 and 2003
- The number of cases in stage 2
- The total number of cases (at stages 1 and 2)

## Results

**Table 1.** Profile of the incidence of new cases of Human African Trypanosomiasis (HAT) in DRC from 2002 to 2003 following the stage of disease progression

A. In 2002
| ENDEMIC AREAS | TPE | ST1 | ST2 | IR-ST1 | IR-ST2 | DELTA |
| --- | --- | --- | --- | --- | --- | --- |
| BAS-CONGO | 146,158 | 183 | 94 | 0.125 | 0.064 | 0.061 |
| BANDUNDU | 587,268 | 2,612 | 1,104 | 0.445 | 0.188 | 0.257 |
| MANIEMA/KATANGA | 99,184 | 329 | 191 | 0.332 | 0.193 | 0.139 |
| KASAI | 179,199 | 704 | 500 | 0.393 | 0.279 | 0.114 |
| **EQUATEUR NORD** | **878,026** | **332** | **642** | **0.038** | **0.073** | **-0.035** |
| EQUATEUR SUD | 80,266 | 27 | 19 | 0.034 | 0.024 | 0.010 |
| TOTAL | 1,970,101 | 4,187 | 2,550 |  |  |  |

B. In 2003
| ENDEMIC AREAS | TPE | ST1 | ST2 | IR-ST1 | IR-ST2 | DELTA |
| --- | --- | --- | --- | --- | --- | --- |
| BAS-CONGO | 180,080 | 162 | 106 | 0.090 | 0.059 | 0.031 |
| BANDUNDU | 758,600 | 2,046 | 1,059 | 0.270 | 0.140 | 0.130 |
| MANIEMA/KATANGA | 123,713 | 292 | 126 | 0.236 | 0.102 | 0.134 |
| KASAI | 205,256 | 561 | 368 | 0.273 | 0.179 | 0.094 |
| EQUATEUR NORD | 949,747 | 203 | 461 | 0.021 | 0.049 | -0.027 |
| EQUATEUR SUD | 94,111 | 18 | 20 | 0.019 | 0.021 | -0.002 |
| TOTAL | 2,311,507 | 3,282 | 2,140 |  |  |  |

Legend: TPE: Total Population Examined; ST1 and ST2: stages 1 and 2; IR-ST1 and IR-ST2: Incidence Rate in stages 1 and 2; DELTA: difference of incidence between stages 1 and 2.

**Table 2.** Evolution from a household to another

**A. In 2002**
| ENDEMICS AREAS | TPT | NC | IR | ENDEMICITY LEVEL |
| --- | --- | --- | --- | --- |
| BAS-CONGO | 146,158 | 760 | 0.52 | Middle |
| **BANDUNDU** | **587,268** | **6,277** | **1.07** | **High** |
| MANIEMA/KATANGA | 99,184 | 590 | 0.59 | Middle |
| **KASAI** | **179,199** | **3,173** | **1.77** | **High** |
| EQUATEUR NORD | 878,026 | 2,436 | 0.28 | Low |
| EQUATEUR SUD | 80,266 | 158 | 0.20 | Low |
| TOTAL | 1,970,101 | 13,394 |  |  |

**B. In 2003**
| ENDEMICS AREAS | TPT | NC | IR | ENDEMICITY LEVEL |
| --- | --- | --- | --- | --- |
| BAS-CONGO | 180,080 | 522 | 0.29 | Middle |
| **BANDUNDU** | **758,600** | **5,384** | **0.71** | **High** |
| MANIEMA/KATANGA | 123,713 | 737 | 0.59 | Middle |
| **KASAI** | **205,256** | **2,606** | **1.27** | **High** |
| EQUATEUR NORD | 949,747 | 1,597 | 0.17 | Low |
| EQUATEUR SUD | 94,111 | 103 | 0.11 | Low |
| TOTAL | 2,311,507 | 10,949 |  |  |

TPT = Total Population Tested; NC = New cases; IR = Incidence Rate

**Table 3.** Rate of confirmed parasitological diagnosis compared with positive CATT from one household to another

**A. In 2002**
| ENDEMICS AREAS | TD+ | CATT + | % CATT+ confirmed |
| --- | --- | --- | --- |
| BAS-CONGO | 246 | 441 | 55 |
| BANDUNDU | 477 | 3,508 | 13 |
| MANIEMA/KATANGA | 158 | 168 | 94 |
| KASAI | 1,021 | 1,286 | 79 |
| EQUATEUR NORD | 1,066 | 2,816 | 37 |
| EQUATEUR SUD | 45 | 51 | 88 |

**B. In 2003**
| ENDEMICS AREAS | TD+ | CATT + | % CATT+ confirmed |
| --- | --- | --- | --- |
| BAS-CONGO | 176 | 386 | 45 |
| BANDUNDU | 570 | 3,397 | 16 |
| MANIEMA/KATANGA | 363 | 286 | 126 |
| KASAI | 811 | 1,046 | 77 |
| EQUATEUR NORD | 647 | 1,089 | 59 |
| EQUATEUR SUD | 46 | 59 | 128 |

TD+ = Thick Drop positive (Parasitological diagnosis); CATT+ = Card Agglutination
Trypanosomiasis Test positive

**Table 4.** Discrepancy rate between parasitological diagnoses and positive CATT from one household to another

**A. In 2002**
| ENDEMICS AREAS | TD+ | CATT + | DISCORDANCE |
| --- | --- | --- | --- |
| BAS-CONGO | 246 | 441 | 55 (discordance) |
| BANDUNDU | 477 | 3,508 | 13 (discordance) |
| MANIEMA/KATANGA | 158 | 168 | 94 (concordance) |
| KASAI | 1,021 | 1,286 | 79 (discordance) |
| EQUATEUR NORD | 1,066 | 2,816 | 37 (discordance) |
| EQUATEUR SUD | 45 | 51 | 88 (concordance) |

**B. In 2003**
| ENDEMICS AREAS | TD+ | CATT + | DISCORDANCE |
| --- | --- | --- | --- |
| BAS-CONGO | 176 | 386 | 45 (discordance) |
| BANDUNDU | 570 | 3,397 | 16 (discordance) |
| MANIEMA/KATANGA | 363 | 286 | 126 (discordance) |
| KASAI | 811 | 1,046 | 77 (discordance) |
| EQUATEUR NORD | 647 | 1,089 | 59 (discordance) |
| EQUATEUR SUD | 46 | 59 | 128 (discordance) |

TD+ = Thick Drop positive (Parasitological diagnosis); CATT+ = Card Agglutination Trypanosomiasis Test positive

**Table 5.** Percentage of stage 2 cases from one period to another

| ENDEMICS AREAS | 2002 | 2003 | DIFFERENCE |
| --- | --- | --- | --- |
| BAS-CONGO | 183 | 162 | 21 |
| BANDUNDU | 2,612 | 2,046 | 566 |
| MANIEMA/KATANGA | 329 | 292 | 37 |
| KASAI | 704 | 561 | 143 |
| EQUATEUR NORD | 332 | 203 | 129 |
| EQUATEUR SUD | 27 | 18 | 9 |
| **TOTAL** | **4,187** | **3,282** | **905** |

## Discussion

### Case discussion for the year 2002

### Impact

In all endemic areas, 1,970,101 people were tested. The national average coverage of the population tested (TPT) represents 16.20% of the exposed population among which 13,853 new cases were detected.

The coverage of the TPT varies from province to province between 4.74% for Maniema / Katanga and 51.11% for Equateur Nord, the overall infection rate is 0.69%.

Compared with the year 2002, the IR decrease from 0.89% to 0.69% and Bandundu took the lead with 46.31% of new cases from all over the country, followed by Kasai 23.41% and Equateur Nord 17.97% with 16.6% new cases. 4 new mobile units were installed in December 2002 including 2 in Bandundu, 2 in Kasaï (**Supportive information 1**).

Of the 13,853 patients reported in 2002, of which 5,636 in the first stage is 52.70% and 7,844 in the 2^nd^ stage is 47.30% with 2.74% unknown stage and the data of Maniema are partial. The unknown or undetermined stage are related to the pregnant woman and the bleeding lumbar puncture (LP) (**Supportive information 2**).

Serum screening or CATT on whole blood is a strategy that the PNLTHA advocates for generalization (3 consecutive years) especially in a household with a high level of endemicity i.e. IR> 1%.

For the year 2002: 1,815,022 people were seen with CATT on whole blood or 92% of the examined population and 9,128 people were confirmed positive in this group is 67% of new cases; 148,019 or 8% of 1,815,022 people had a positive result at CATT (**Supportive information 3**). Cases diagnosed by other methods in Maniema and Equateur Sud do not have a known method of diagnosis.

PNLTHA recommends the use of Pentamidine and Bayer for the treatment of patients in the first phase, Melarsoprol for the second phase while DFMO and LAMPIT are reserved as relay drugs (**Supportive information 4**).

The lumbar puncture performed was 28,253 or 58.6% of the controls performed. PLC abnormal 7,187 is 25.4% i.e. the elements are not the normal limits i.e. <5 / mm^3^, this explains the difficulty the PNLTHA have for the post-therapeutic follow-up. Most patients, once out, if they feel better do not come back. An effort should be made to identify relapse cases (**Supportive information 8**).

### Extent of the problem at the national level

The population at risk is estimated at 12,600,000 people representing more than 20% of the population of the Democratic Republic of the Congo and 23% of the total population at risk of Africa which is estimated by the WHO to 60 million.

To date, DRC continues to bear the heaviest burden of gambiense HAT, having reported 84 % of all African cases in 2012 (i.e., 5,968/7,106). Therefore, achieving the international goal of gambiense HAT elimination [3], will depend to a large extent on the progress that DRC will be able to make [4] in ridding gambiense HAT following the Yerevan resolution on neglected tropical diseases (https://www.who.int/neglected_diseases/news/OIF-commit-to-strengthening-the-fight-against-NTDs/en/).

The monthly average of new cases in the country was 1,154 in 2002, the at-risk population covered by specialized services and primary health care in 2002 represents 16% of the total population at risk, or 2,040,927 persons, of which 88% is 1,815,022 were tested with CATT, which represents approximately 14.4% of the exposed population. Of these 88% came out 9,128 new cases. This coverage varies from province to province. It ranges from 4.74% in Maniema to 51% in Equateur Nord.

### Extent of the problem at the provincial level

The following provinces reported 87% of new cases in 2002: 46.3% Bandundu; 23.4% in Kasaï and 17.9% in Equateur Nord.

### Case discussion for the year 2003

### Impact

In all endemic areas, 2,405,991 people were examined. The national average coverage of the TPT represents 19.10% of the exposed population. The number of new Human African Trypanosomiasis (HAT) cases is 11,481 (**Supportive information 9**). Among the 11,481 new HAT patients declared:

- 4,339 or 37.8% at the 1^st^ stage (lymphatic-blood)

- 6,913 or 60.22% at the 2^nd^ stage (meningoencephalopathic)

- 229 or about 1.98% whose stadium diagnosis was not made because of:

* refusal of the lumbar puncture or haemorrhagic not repeated

* The gravid status of the patient

- 5,786 new cases were detected by specialized units

- 5,695 new cases were detected by fixed structures (CDTC and health facilities) (**Supportive information 10**).

For the year 2003; 2,166,835 people were seen with CATT test on whole blood or 94% of the population examined (**Supportive information 11**).

Cases diagnosed by other methods in Maniema and Equateur Sud have no known diagnostic method (**Supportive information 12**).

PNLTHA recommends the use of Pentamidine and Bayer for the treatment of patients in the first phase, Melarsoprol for the second phase while the DFMO and Lampit are reserved as relay drugs. Arsobal remains the most used drug (**Supportive information 13**).

### Extent of the problem at national level

The monthly average of new cases in Democratic Republic of the Congo was 957 in 2003. (**Supportive information 14**)

TPE coverage varies from province to province the other between 8.17% for Equateur Sud and 52.69% for Equateur Nord.

The overall infection rate decreased from 0.89% to 0.48% in 2003. 4 new mobile units (MU) were installed in December 2002, including 2 in Bandundu and contributed to the work during 2003. Thanks to the funding of the WHO for the 3 MU and Belgian Technical Cooperation for 1 MU.

The eastern province provided after ISANGI survey 251 NC out of 4,140 people tested with a detection rate of 6.06 highlighting the severity of the endemic and calling for an organization to fight the Human African Trypanosomiasis in the Democratic Republic of the Congo following the resolution of 2018 French Summit Committee in Yerevan (Armenia) on the Neglected Tropical Diseases (https://www.who.int/neglected_diseases/news/OIF-commit-to-strengthening-the-fight-against-NTDs/en/).

### Extent of the problem at the provincial level

Bandundu is in the lead with 47% of new cases from across the country, followed by Kasai 22.6% and Equateur Nord 14.80%

### Case discussion for 2002 and 2003

Table 1 shows that the infection rate decreased from 2002 to 2003 in all households except Equateur Nord, while Table 2 reports the severity of sleeping sickness in the Bandundu provinces. and from Kasai, Table 3 extended the discrepancy between the parasitological test (thick drop) and the serological test and raised the problem of CATT sensitivity; finally, Table 4 showed that the number of cases detected in stage 2 has decreased (4,187 cases detected in 2002 against 3,282 cases detected in 2003, a difference of 905 cases), hence we encourage the PNLTHA to continue in this direction of the reduction of HAT-related morbidity.

The total number of new cases of Human African Trypanosomiasis (HAT) in the Democratic Republic of the Congo declined sharply from 17,322 in 2001 to 13,853 in 2002, a decrease related to the number of new cases in Equateur Nord. In the provinces of Bandundu and Kasaï, however, the problem of Human African Trypanosomiasis (HAT) remains important.

We see a growing evolution of the population examined from year to year. This would be even higher if the wars had not caused the slowdown in active screening because of insecurity in the combat zones. The new cases detected show the effort deployed by the heroes of the shadows, and once the peace is restored, the work’s breadth (**Supportive information 7 and 14**).

Serum screening or CATT on whole blood is a strategy that the PNLTHA advocates for generalization (3 consecutive years) especially in a household with a high level of endemicity i.e. IR> 1%.

## Conclusion

In short, we say that the actual number of people already infected is probably higher than the new cases reported. We are still far from the situation of 1958/60 when there was 1 new case declared by 10,000 people examined (i.e. 1,100 new cases out of 13,000,000 people seen). Therefore, the Congolese government must make long-term financial commitments to ensure the continuity of HAT control activities.

## SUPPORTIVE INFORMATION

### 1. RESULTS OF HAT ACTIVITIES IN 2002 BY ENDEMIC ZONE

**S 1:** Distribution by Endemic Area of Total Registered Population (TRP), Total Enrolled Population (TEP) of New Cases (NC) and Infection Rate (IR) in 2002

| ENDEMICS AREAS | EXPOSED POP | Nbr HZ | HZ HAT | Nbr MU |  | TPC | TPE | % cover | TP | NC | IR | Level of Endemicity |
| --- | --- | --- | --- | --- | --- | --- | --- | --- | --- | --- | --- | --- |
|  |  |  |  | 1<sup>st</sup> S e m | 2<sup>nd</sup> S e m |  |  |  |  |  |  |  |
| BAS-CONGO | 1,700,000 | 27 | 19 | 5 | 5 | 228,431 | 146,158 | 13.44 | 63.98 | 760 | 0.52 | Middle |
| BANDUNDU | 3,800,000 | 38 | 31 | 12 | 14 | 767,966 | 587,268 | 20.21 | 76.47 | 6,277 | 1.07 | Severe |
| MANIEMA /KATANGA | 1,200,000 | 81 | 16 | 4 | 4 | 81,994 | 99,184 | 6.83 | 120.96 | 590 | 0.59 | Middle |
| KASAI | 2,000,000 | 58 | 35 | 5 | 7 | 294,764 | 179,199 | 14.74 | 60.79 | 3,173 | 1.77 | Severe |
| EQUATEUR NORD | 2,000,000 | 20 | 15 | 9 | 9 | 1,022,219 | 878,026 | 51.11 | 85.89 | 2,436 | 0.28 | Low |
| EQUATEUR SUD | 1,000,000 | 13 | 6 | 2 | 2 | 77,743 | 80,266 | 7.77 | 103.25 | 158 | 0.20 | Middle |
| KINSHASA | 600,000 | 22 | 8 | 2 | 2 | 80,644 | 70,826 | 13.44 | 87.83 | 459 | 0.65 | Middle |
| PROVINCE ORIENTALE | 300,000 | 47 | 3 | NA | NA | NA | NA | NA | NA | NA | NA | NA |
| TOTALS | 12,600,000 | 306 | 133 | 39 | 43 | 2,553,761 | 2,040,927 | **16.20** | 79.92 | 13,853 | 0.68 | Middle |

Legend: HZ = Health Zone; HZ HAT = Health Zone at risk of trypanosomiasis; MU = Mobile Unit; TPC = Total Population Counted; NA=Not Available or missing data
PTE = Total Population Examined; NC = new cases; IR = Infection Rate

**S 2:** Distribution by endemic area of new cases recorded according to the screening circumstances and stage of the disease in the DRC in 2002

| Circumstances of screening | TPT | TPT AS | TPT PS | ACTIVE SCREENING |  |  |  | PASSIVE SCREENING |  |  |  | TOT NC |
| --- | --- | --- | --- | --- | --- | --- | --- | --- | --- | --- | --- | --- |
|  |  |  |  | 1<sup>st</sup> stage | 2<sup>nd</sup> stage | Unk own | Total | 1<sup>st</sup> stage | 2<sup>nd</sup> stage | Unk own | Total |  |
| Endemics areas |  |  |  |  |  |  |  |  |  |  |  |  |
| BAS-CONGO | 146,158 | 142,321 | 3,837 | 183 | 94 | NA | 277 | 175 | 308 | NA | 483 | 760 |
| BANDUNDU | 587,268 | 551,568 | 35,700 | 2,612 | 1,104 | 163 | 3,879 | 605 | 1,783 | 10 | 2,398 | 6,277 |
| MANIEMA/KATANGA | 99,184 | 97,929 | 1,255 | 329 | 191 | 7 | 527 | 17 | 46 | NA | 63 | 590 |
| KASAI | 179,199 | 165,532 | 13,667 | 704 | 500 | 88 | 1,292 | 251 | 1,615 | 15 | 1,881 | 3,173 |
| EQUATEUR NORD | 878,026 | 822,283 | 55,743 | 332 | 642 | 50 | 1,024 | 211 | 1,190 | 11 | 1,412 | 2,436 |
| EQUATEUR SUD | 80,266 | 79,213 | 1,053 | 27 | 19 | 1 | 47 | 11 | 100 | NA | 111 | 158 |
| KINSHASA | 70,826 | 67,170 | 3,656 | 141 | 88 | 26 | 255 | 38 | 164 | 2 | 204 | 459 |
| PROVINCE ORIENTALE | 0 | 0 | 0 | NA | NA | NA | 0 | NA | NA | NA | 0 | 0 |
| TOTALS | 2,040,927 | 1,926,016 | 114,911 | 4,328 | 2,638 | 335 | 7,301 | 1,308 | 5,206 | 36 | 6,552 | 13,853 |
| PERCENTAGE (%) |  |  |  |  |  |  | 52.70 |  |  |  | 47.30 |  |

Legend: TPT = Total Population Tested; TPT AS = Total Population Tested by Active Screening; TPT PS = total population tested by passive screening; NA=Not Available or missing data, NC = new case

**S 3:**
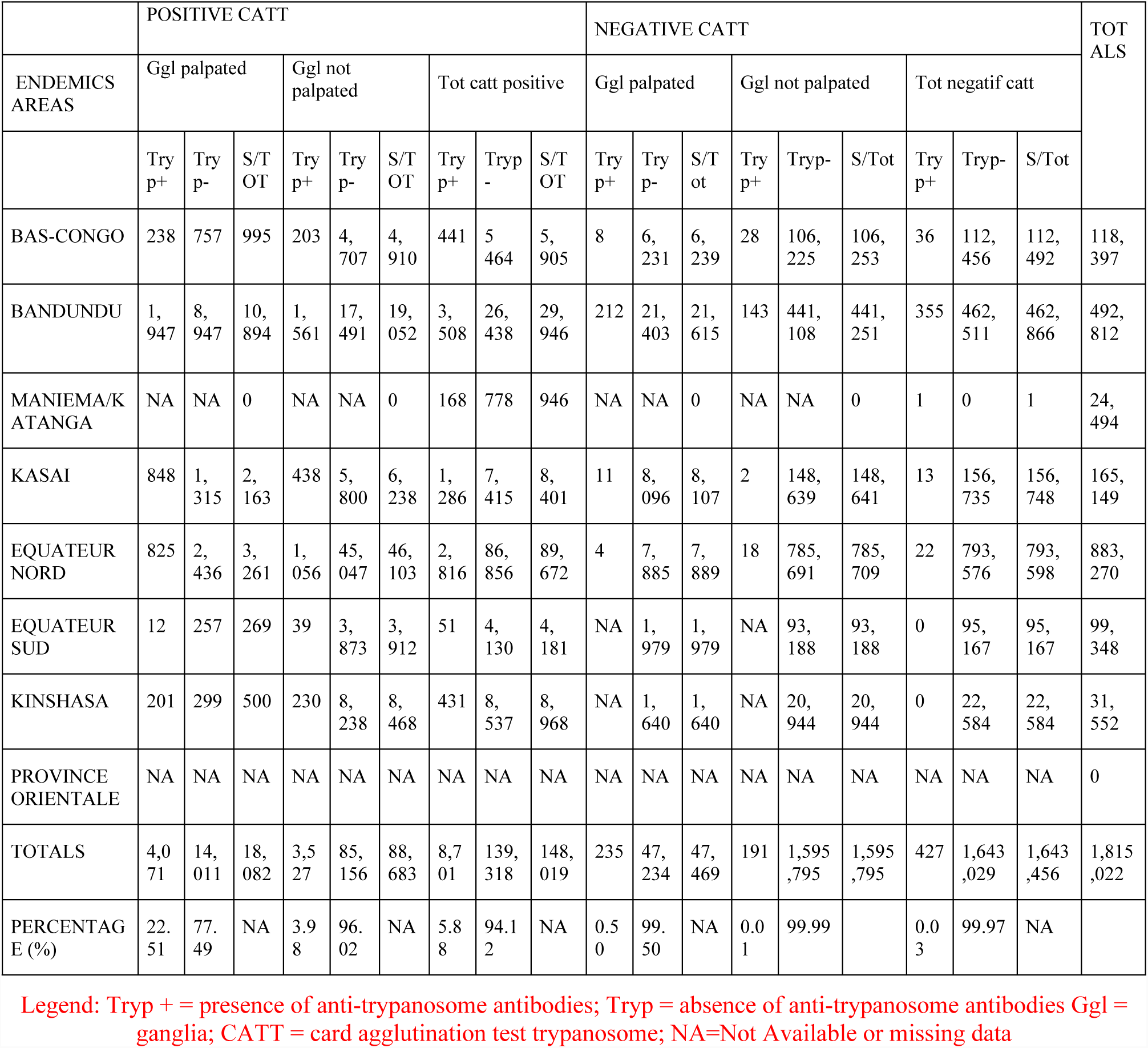
CATT test results in relation to ganglia in 2002

**S4:** Distribution by endemic area of new cases detected according to the mode of diagnosis in the DRC in 2002

| Diagnosis mode | GP | FB | TD | TBC | CSF | CLIN | OTHERS WOO + DC<br>LC | TOTAL |
| --- | --- | --- | --- | --- | --- | --- | --- | --- |
| Endemic areas |  |  |  |  |  |  |  |  |
| BAS-CONGO | 419 | 51 | 246 | NA | 1 | NA | 43 | 760 |
| BANDUNDU | 3,158 | 2,018 | 477 | 23 | 553 | 1 | 47 | 6,277 |
| MANIEMA/KATANGA | 235 | 153 | 158 | NA | 23 | 21 | 0 | 590 |
| KASAI | 1,782 | 54 | 1,021 | 2 | 291 | 17 | 6 | 3,173 |
| EQUATEUR NORD | 875 | 174 | 1,066 | NA | 267 | 48 | 6 | 2,436 |
| EQUATEUR SUD | 27 | 21 | 45 | 1 | 11 | NA | 53 | 158 |
| KINSHASA | 176 | 21 | 174 | 6 | 61 | 1 | 20 | 459 |
| PROVINCE<br>ORIENTALE | NA | NA | NA | NA | NA | NA | NA | 0 |
| TOTALS | 6,672 | 2,492 | 3,187 | 32 | 1,207 | 88 | 175 | 13,853 |
| DETECTION RATE | 48.16 | 17.99 | 23.01 | 0.23 | 8.71 | 0.64 | 1.26 |  |

Legend: GP = ganglionic puncture; FB = fresh blood; TD = thick drop (parasitology diagnosis); TBC = triple blood centrifugation; CSF = cerebrospinal fluid, NA=Not Available or missing data

**S5:**
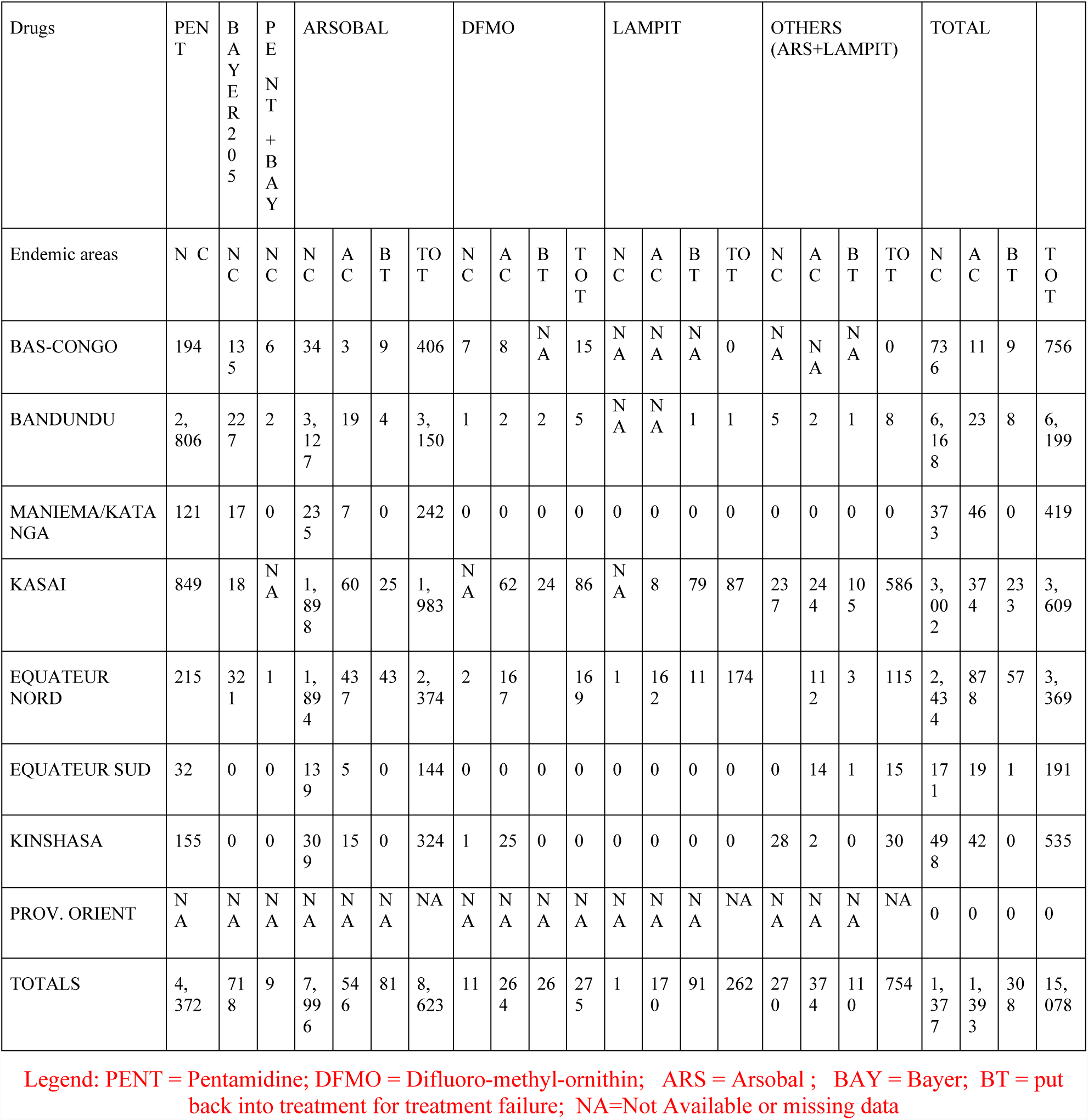
distribution by endemic area of patients treated according to the therapeutic regime in 2002

**S6:** Distribution of the cases following the control structures in 2002

**S6: Distribution of the cases following the control structures in 2002**
| Endemic areas | BAS-CONGO | BANDUNDU | MANIEMA/<br>KATANGA | KASAI | EQUATEUR<br>NORD | EQUATEUR<br>SUD | KINSHASA | PROVINCE<br>ORIENTALE | TOTALS |
| --- | --- | --- | --- | --- | --- | --- | --- | --- | --- |
| Control Structures |  |  |  |  |  |  |  |  |  |
| MU | 316 | 3,936 | 463 | 1,267 | 1,024 | 54 | 257 | 0 | 7,317 |
| CDTC | 374 | 978 | 109 | 951 | 1,402 | 183 | 191 | 0 | 4,108 |
| FS | 70 | 1,363 | 18 | 955 | 10 | 1 | 11 | 0 | 2,428 |
| TOTAL | 760 | 6,277 | 590 | 3,173 | 2,436 | 158 | 459 | 0 | 13,853 |

Legend: MU= Mobile Unit

**S 7:** Evolution of HAT in the last six years in the DRC; TPE / 1NC report and monthly average of NC by year

**S 7: Evolution of HAT in the last six years in the DRC; TPE / INC report and monthly average of NC by year**
| YEAR | 1996 | 1997 | 1998 | 1999 | 2000 | 2001 | 2002 |
| --- | --- | --- | --- | --- | --- | --- | --- |
| DATA COLLECTED |  |  |  |  |  |  |  |
| TPT | 1,297,778 | 1,168,404 | 1,472,674 | 1,298,933 | 1,605,816 | 2,028,238 | 2,040,927 |
| NC | 19,485 | 25,094 | 26,318 | 18,684 | 16,975 | 17,322 | 13,853 |
| Infection rate | 1.50 | 2.15 | 1.79 | 1.44 | 1.06 | 0.85 | 0.68 |
| Monthly average New cases | 1,624 | 2,091 | 2,193 | 1,557 | 1,415 | 1,444 | 1,154 |

**Legend: TPT = Total Population Tested; NC = new cases; IR=Infection Rate**

**S 8:** Lumbar punctures for diagnosis and control in 2002

**Legend: TPT = Total Population Tested; NC = new cases; IR=Infection Rate** **S 8: Lumbar punctures for diagnosis and control in 2002**
| LP |  |  |  |  | 2002 |  |  |  |
| --- | --- | --- | --- | --- | --- | --- | --- | --- |
| Endemics Areas | NC 2000 | NC 2001 | NC 2002 | Prevision LPC 2002 | PLDA-DB | TLPC | TPLCN | TLPAC |
| BAS-CONGO | 816 | 736 | 760 | 2312 | 760 | 1,926 | 1,664 | 262 |
| BANDUNDU | 5,588 | 7,190 | 6,277 | 19,055 | 6,104 | 5,289 | 5,232 | 57 |
| MANIEMA/KATANGA | 323 | 711 | 590 | 1,624 | 286 | NA | NA | 0 |
| KASAI | 2,810 | 2,905 | 3,173 | 8,888 | 3,070 | 3,854 | 1,697 | 2,157 |
| EQUATEUR NORD | 6,673 | 4,990 | 2,436 | 14,099 | 2,375 | 15,988 | 11,769 | 4,219 |
| EQUATEUR SUD | 139 | 106 | 158 | 403 | 157 | 565 | 173 | 3,992 |
| KINSHASA | NA | NA | NA | NA | NA | NA | NA | NA |
| PROVINCE ORIENTALE | 16,975 | 17,319 | 1,385,553 | 48,147 | 13,183 | 28,253 | 21,066 | 7,187 |
| TOTALS |  |  |  |  |  |  |  |  |

Legend: LP = lumbar puncture; NC = new case; PLC LPC= lumbar puncture control; TLPC = Total Lumbar Puncture Control; TLPNC = total lumbar puncture normal control; TLPAC= total lumbar puncture of abnormal control, NA=Not Available or missing data

### 2. RESULTS OF 2003 THA CONTROL ACTIVITIES BY ENDEMIC ZONE

**S 9:**
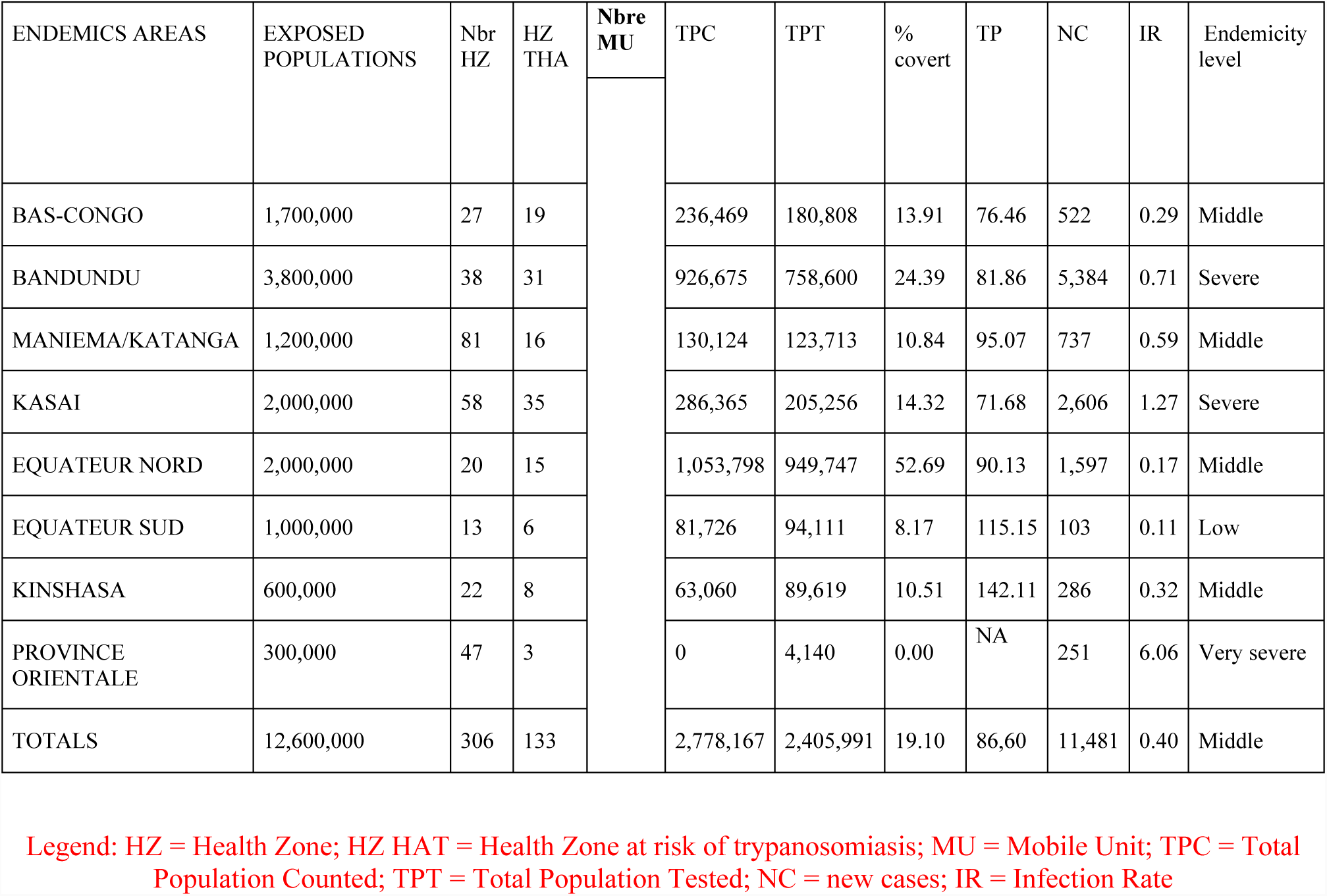
Distribution by Endemic Area of Total Registered Population (TRP), Total Enrolled Population (TEP) of New Cases (NC) and Infection Rate (IR) in 2003

| ENDEMIC AREAS | EXPOSED POPULATIONS | Nbr HZ | HZ THA | Nbre MU | TPC | TPT | % covert | TP | NC | IR | Endemicity level |
| --- | --- | --- | --- | --- | --- | --- | --- | --- | --- | --- | --- |
| BAS-CONGO | 1,700,000 | 27 | 19 |  | 236,469 | 180,808 | 13.91 | 76.46 | 522 | 0.29 | Middle |
| BANDUNDU | 3,800,000 | 38 | 31 |  | 926,675 | 758,600 | 24.39 | 81.86 | 5,384 | 0.71 | Severe |
| MANIEMA/KATANGA | 1,200,000 | 81 | 16 |  | 130,124 | 123,713 | 10.84 | 95.07 | 737 | 0.59 | Middle |
| KASAI | 2,000,000 | 58 | 35 |  | 286,365 | 205,256 | 14.32 | 71.68 | 2,606 | 1.27 | Severe |
| EQUATEUR NORD | 2,000,000 | 20 | 15 |  | 1,053,798 | 949,747 | 52.69 | 90.13 | 1,597 | 0.17 | Middle |
| EQUATEUR SUD | 1,000,000 | 13 | 6 |  | 81,726 | 94,111 | 8.17 | 115.15 | 103 | 0.11 | Low |
| KINSHASA | 600,000 | 22 | 8 |  | 63,060 | 89,619 | 10.51 | 142.11 | 286 | 0.32 | Middle |
| PROVINCE ORIENTALE | 300,000 | 47 | 3 |  | 0 | 4,140 | 0.00 | NA | 251 | 6.06 | Very severe |
| TOTALS | 12,600,000 | 306 | 133 |  | 2,778,167 | 2,405,991 | 19.10 | 86.60 | 11,481 | 0.40 | Middle |

Legend: HZ = Health Zone; HZ HAT = Health Zone at risk of trypanosomiasis; MU = Mobile Unit; TPC = Total Population Counted; TPT = Total Population Tested; NC = new cases; IR = Infection Rate

**S 10:** Distribution by endemic area of new cases recorded according to the screening circumstances and stage of the disease in the DRC in 2002

| Circumstances of screening | ACTIVE SCREENING |  |  |  | PASSIVE SCREENING |  |  |  | TOT<br>NC |
| --- | --- | --- | --- | --- | --- | --- | --- | --- | --- |
|  | 1<sup>st</sup><br>stage | 2<sup>nd</sup><br>stage | Unkown | Total | 1<sup>st</sup><br>stage | 2<sup>nd</sup><br>stage | Unkown | Total |  |
| Endemics<br>Areas |  |  |  |  |  |  |  |  |  |
| BAS-CONGO | 162 | 106 | 2 | 270 | 67 | 185 | 0 | 252 | 522 |
| BANDUNDU | 2,046 | 1,059 | 77 | 3,182 | 547 | 1,646 | 9 | 2,202 | 5,384 |
| MANIEMA/KATANGA | 292 | 126 | 25 | 443 | 58 | 231 | 0 | 289 | 732 |
| KASAI | 561 | 368 | 41 | 970 | 157 | 1465 | 14 | 1,636 | 2,606 |
| EQUATEUR NORD | 203 | 461 | 16 | 680 | 75 | 835 | 7 | 917 | 1,597 |
| EQUATEUR SUD | 18 | 20 | 2 | 40 | 12 | 50 | 1 | 63 | 103 |
| KINSHASA | 43 | 40 | 22 | 105 | 18 | 163 | 0 | 181 | 286 |
| PROVINCE ORIENTALE | 34 | 54 | 8 | 96 | 46 | 104 | 5 | 155 | 251 |
| TOTALS | 3,359 | 2,234 | 193 | 5,786 | 980 | 4,679 | 36 | 5,695 | 11,481 |
| PERCENTAGE (%) |  |  |  | 50.40 |  |  |  | 49.60 |  |

Legend: NC = new cases

**S 11:**
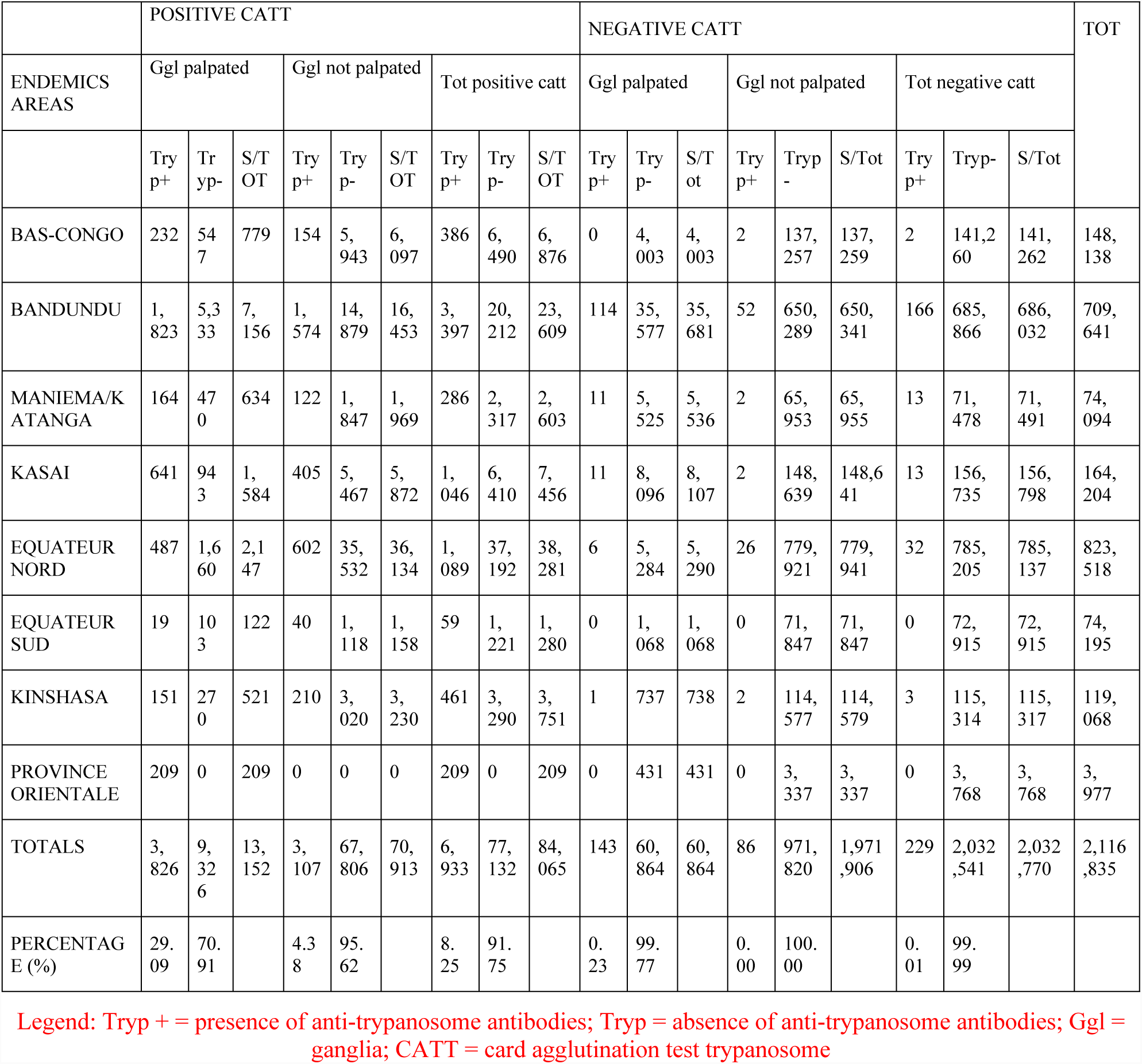
CATT test results for ganglia in 2003

**S 11: CATT test results for ganglia in 2003**
|  | POSITIVE CATT |  |  |  |  |  |  |  |  | NEGATIVE CATT |  |  |  |  |  |  |  |  | TOT |
| --- | --- | --- | --- | --- | --- | --- | --- | --- | --- | --- | --- | --- | --- | --- | --- | --- | --- | --- | --- |
| ENDEMIC AREAS | Ggl palpated |  |  | Ggl not palpated |  |  | Tot positive catt |  |  | Ggl palpated |  |  | Ggl not palpated |  |  | Tot negative catt |  |  |  |
|  | Try p+ | Try yp- | S/T OT | Try p+ | Try p- | S/T OT | Try p+ | Try p- | S/T OT | Try p+ | Try p- | S/T ot | Try p+ | Tryp - | S/Tot | Try p+ | Tryp- | S/Tot |  |
| BAS-CONGO | 232 | 547 | 779 | 154 | 5,943 | 6,097 | 386 | 6,490 | 6,876 | 0 | 4,003 | 4,003 | 2 | 137,257 | 137,259 | 2 | 141,260 | 141,262 | 148,138 |
| BANDUNDU | 1,823 | 5,333 | 7,156 | 1,574 | 14,879 | 16,453 | 3,397 | 20,212 | 23,609 | 114 | 35,577 | 35,681 | 52 | 650,289 | 650,341 | 166 | 685,866 | 686,032 | 709,641 |
| MANIEMA/KATANGA | 164 | 470 | 634 | 122 | 1,847 | 1,969 | 286 | 2,317 | 2,603 | 11 | 5,525 | 5,536 | 2 | 65,953 | 65,955 | 13 | 71,478 | 71,491 | 74,094 |
| KASAI | 641 | 943 | 1,584 | 405 | 5,467 | 5,872 | 1,046 | 6,410 | 7,456 | 11 | 8,096 | 8,107 | 2 | 148,639 | 148,641 | 13 | 156,735 | 156,798 | 164,204 |
| EQUATEUR NORD | 487 | 1,660 | 2,147 | 602 | 35,532 | 36,134 | 1,089 | 37,192 | 38,281 | 6 | 5,284 | 5,290 | 26 | 779,921 | 779,941 | 32 | 785,205 | 785,137 | 823,518 |
| EQUATEUR SUD | 19 | 103 | 122 | 40 | 1,118 | 1,158 | 59 | 1,221 | 1,280 | 0 | 1,068 | 1,068 | 0 | 71,847 | 71,847 | 0 | 72,915 | 72,915 | 74,195 |
| KINSHASA | 151 | 270 | 521 | 210 | 3,020 | 3,230 | 461 | 3,290 | 3,751 | 1 | 737 | 738 | 2 | 114,577 | 114,579 | 3 | 115,314 | 115,317 | 119,068 |
| PROVINCE ORIENTALE | 209 | 0 | 209 | 0 | 0 | 0 | 209 | 0 | 209 | 0 | 431 | 431 | 0 | 3,337 | 3,337 | 0 | 3,768 | 3,768 | 3,977 |
| TOTALS | 3,826 | 9,326 | 13,152 | 3,107 | 67,806 | 70,913 | 6,933 | 77,132 | 84,065 | 143 | 60,864 | 60,864 | 86 | 971,820 | 1,971,906 | 229 | 2,032,541 | 2,032,770 | 2,116,835 |
| PERCENTAGE (%) | 29.09 | 70.91 |  | 4.38 | 95.62 |  | 8.25 | 91.75 |  | 0.23 | 99.77 |  | 0.00 | 100.00 |  | 0.01 | 99.99 |  |  |

Legend: Tryp + = presence of anti-trypanosome antibodies; Tryp - = absence of anti-trypanosome antibodies; Ggl = ganglia; CATT = card agglutination test trypanosome

**S 12:** Distribution by endemic area of new cases detected according to the diagnostic method in the DRC in 2003

| Diagnosis mode | GP | FB | TD | TBC | CSF | CLIN | OTHERS WOO + DC LC | TOTAL |
| --- | --- | --- | --- | --- | --- | --- | --- | --- |
| Endemic areas |  |  |  |  |  |  |  |  |
| BAS-CONGO | 276 | 26 | 176 | 0 | 44 | 0 | 0 | 522 |
| BANDUNDU | 2,679 | 1,455 | 570 | 519 | 21 | 0 | 140 | 5,384 |
| MANIEMA/KATANGA | 240 | 63 | 363 | 65 | 0 | 1 | 0 | 732 |
| KASAI | 1,317 | 38 | 811 | 0 | 393 | 18 | 29 | 2,606 |
| EQUATEUR NORD | 586 | 94 | 647 | 0 | 208 | 55 | 7 | 1,595 |
| EQUATEUR SUD | 36 | 7 | 46 | 0 | 14 | 0 | 0 | 103 |
| KINSHASA | 114 | 13 | 65 | 5 | 43 | 0 | 46 | 286 |
| PROVINCE ORIENTALE | 125 | 3 | 100 | 0 | 21 | 0 | 2 | 251 |
| TOTALS | 5,373 | 1,699 | 2,778 | 589 | 744 | 74 | 224 | 11,881 |
| DETECTION RATE | 46.80 | 14.80 | 24.20 | 5.13 | 6.48 | 0.64 | 1.95 |  |

Legend: GP = ganglionic puncture; FB = fresh blood; TD = thick drop; TBC = triple blood centrifugation; CSF = cerebrospinal fluid

**S 13:**
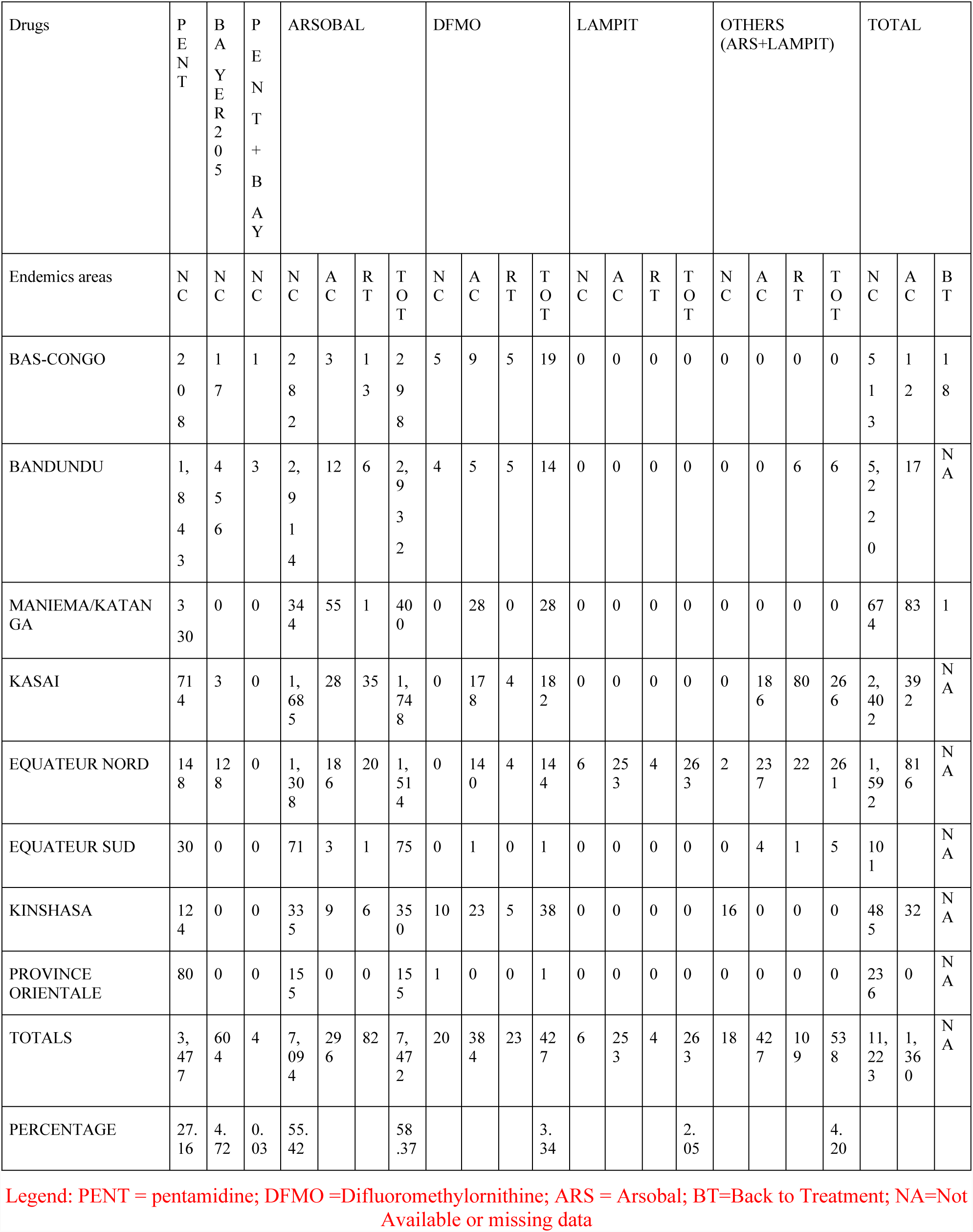
distribution by endemic area of patients treated according to the therapeutic regime in 2003

| Drugs | P<br>E<br>N<br>T | B<br>A<br>Y<br>E<br>R<br>2<br>0<br>5 | P<br>E<br>N<br>T<br>+<br>B<br>A<br>Y | ARSOBAL |  |  |  | DFMO |  |  |  | LAMPIT |  |  |  | OTHERS<br>(ARS+LAMPIT) |  |  |  | TOTAL |  |  |
| --- | --- | --- | --- | --- | --- | --- | --- | --- | --- | --- | --- | --- | --- | --- | --- | --- | --- | --- | --- | --- | --- | --- |
| Endemics areas | N<br>C | N<br>C | N<br>C | N<br>C | A<br>C | R<br>T | T<br>O<br>T | N<br>C | A<br>C | R<br>T | T<br>O<br>T | N<br>C | A<br>C | R<br>T | T<br>O<br>T | N<br>C | A<br>C | R<br>T | T<br>O<br>T | N<br>C | A<br>C | B<br>T |
| BAS-CONGO | 2<br>0<br>8 | 1<br>7 | 1 | 2<br>8<br>2 | 3 | 1<br>3 | 2<br>9<br>8 | 5 | 9 | 5 | 19 | 0 | 0 | 0 | 0 | 0 | 0 | 0 | 0 | 5<br>1<br>3 | 1<br>2 | 1<br>8 |
| BANDUNDU | 1,<br>8<br>4<br>3 | 4<br>5<br>6 | 3 | 2,<br>9<br>1<br>4 | 12 | 6 | 2,<br>9<br>3<br>2 | 4 | 5 | 5 | 14 | 0 | 0 | 0 | 0 | 0 | 0 | 6 | 6 | 5,<br>2<br>2<br>0 | 17 | N<br>A |
| MANIEMA/KATAN<br>GA | 3<br>30 | 0 | 0 | 34<br>4 | 55 | 1 | 40<br>0 | 0 | 28 | 0 | 28 | 0 | 0 | 0 | 0 | 0 | 0 | 0 | 0 | 67<br>4 | 83 | 1 |
| KASAI | 71<br>4 | 3 | 0 | 1,<br>68<br>5 | 28 | 35 | 1,<br>74<br>8 | 0 | 17<br>8 | 4 | 18<br>2 | 0 | 0 | 0 | 0 | 0 | 18<br>6 | 80 | 26<br>6 | 2,<br>40<br>2 | 39<br>2 | N<br>A |
| EQUATEUR NORD | 14<br>8 | 12<br>8 | 0 | 1,<br>30<br>8 | 18<br>6 | 20 | 1,<br>51<br>4 | 0 | 14<br>0 | 4 | 14<br>4 | 6 | 25<br>3 | 4 | 26<br>3 | 2 | 23<br>7 | 22 | 26<br>1 | 1,<br>59<br>2 | 81<br>6 | N<br>A |
| EQUATEUR SUD | 30 | 0 | 0 | 71 | 3 | 1 | 75 | 0 | 1 | 0 | 1 | 0 | 0 | 0 | 0 | 0 | 4 | 1 | 5 | 10<br>1 |  | N<br>A |
| KINSHASA | 12<br>4 | 0 | 0 | 33<br>5 | 9 | 6 | 35<br>0 | 10 | 23 | 5 | 38 | 0 | 0 | 0 | 0 | 16 | 0 | 0 | 0 | 48<br>5 | 32 | N<br>A |
| PROVINCE<br>ORIENTALE | 80 | 0 | 0 | 15<br>5 | 0 | 0 | 15<br>5 | 1 | 0 | 0 | 1 | 0 | 0 | 0 | 0 | 0 | 0 | 0 | 0 | 23<br>6 | 0 | N<br>A |
| TOTALS | 3,<br>47<br>7 | 60<br>4 | 4 | 7,<br>09<br>4 | 29<br>6 | 82 | 7,<br>47<br>2 | 20 | 38<br>4 | 23 | 42<br>7 | 6 | 25<br>3 | 4 | 26<br>3 | 18 | 42<br>7 | 10<br>9 | 53<br>8 | 11,<br>22<br>3 | 1,<br>36<br>0 | N<br>A |
| PERCENTAGE | 27.<br>16 | 4.<br>72 | 0.<br>03 | 55.<br>42 |  |  | 58<br>.37 |  |  |  | 3.<br>34 |  |  |  | 2.<br>05 |  |  |  | 4.<br>20 |  |  |  |

Legend: PENT = pentamidine; DFMO = Difluoromethylornithine; ARS = Arsobal; BT=Back to Treatment; NA=Not Available or missing data

**S 14:** Evolution of HAT in the last eight years in the DRC; PTE / NC ratio and monthly average of NC by year

| YEAR | 1996 | 1997 | 1998 | 1999 | 2000 | 2001 | 2002 | 2003 |
| --- | --- | --- | --- | --- | --- | --- | --- | --- |
| DATA COLLECTED |  |  |  |  |  |  |  |  |
| TPT | 1,297,778 | 1,168,404 | 1,472,674 | 1,298,933 | 1,605,816 | 2,028,238 | 1,973,885 | 2,400,673 |
| NC | 19,485 | 25,094 | 26,318 | 18,684 | 16,975 | 17,322 | 13,553 | 11,481 |
| IR | 1.50 | 2.15 | 1.79 | 1.44 | 1.06 | 0.85 | 0.69 | 0.48 |
| MANC | 1,624 | 2,091 | 2,193 | 1,557 | 1,415 | 1,444 | 1,129 | 957 |

**Legend: TPT = Total Population Tested; NC = New Case; IR= Infection Rate; MANC= Monthly average New cases**

**S15:**
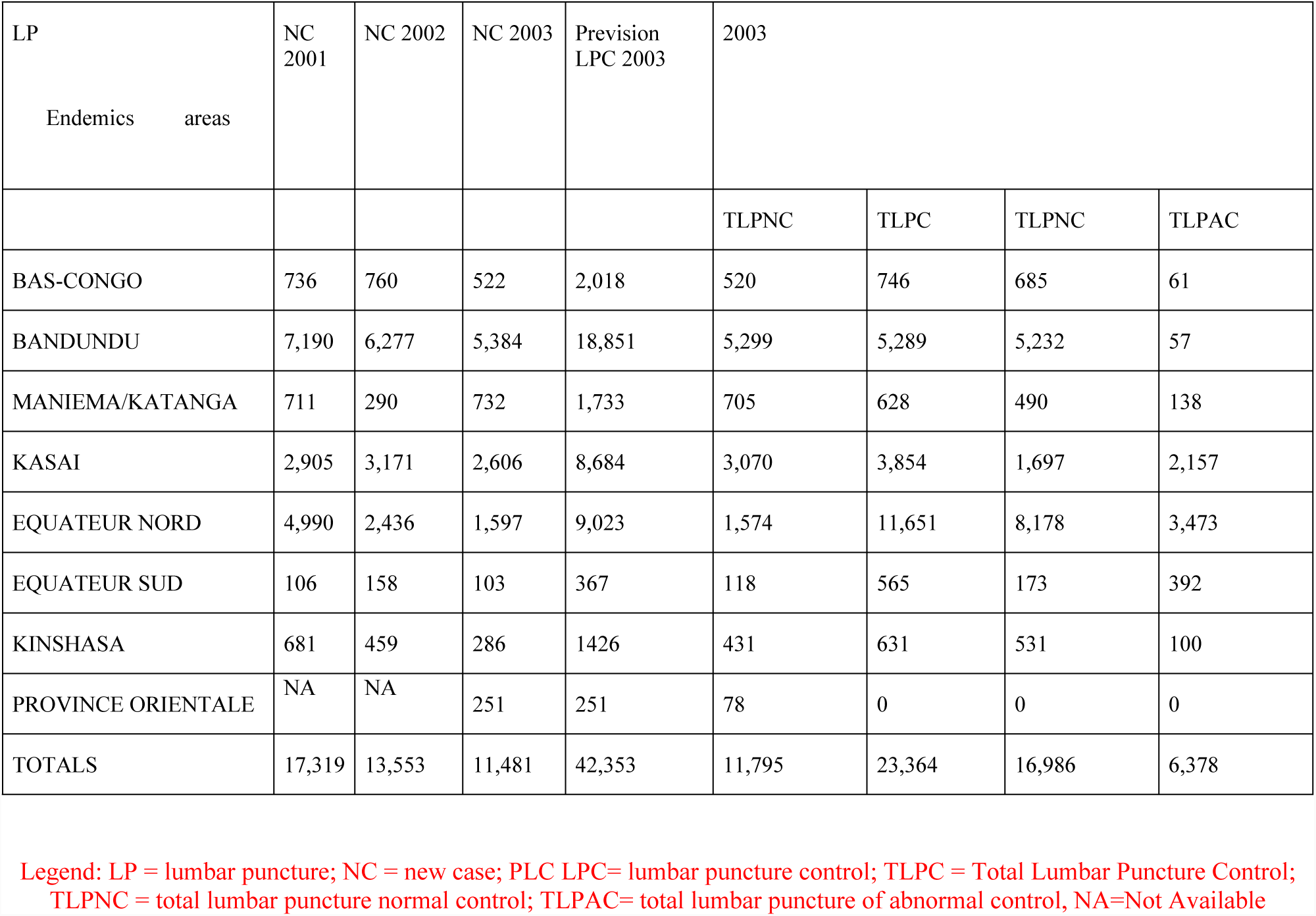
The lumbar punctures of diagnosis and control carried out in 2003

## References

1. Kennedy P.G. Clinical features, diagnosis, and treatment of human African trypanosomiasis (sleeping sickness). Lancet Neurol., 2013, 12, 186–194.

2. Lutumba P, Robays J, Miaka mia Bilenge C, et al. Trypanosomiasis control, Democratic Republic of Congo, 1993-2003. Emerg Infect Dis. 2005;11(9):1382–8.

3. WHO. Report of a WHO meeting on elimination of African trypanosomiasis (Trypanosoma brucei gambiense). Geneva: World Health Organization; 2013. p. 2013. Available from: http://apps.who.int/iris/bitstream/10665/79689/1/WHO_HTM_NTD_IDM_2013.4_eng.pdf?ua=1

4. Lumbala C, Simarro PP, Cecchi G, et al. Human African trypanosomiasis in the Democratic Republic of the Congo: disease distribution and risk. Int J Health Geogr. 2015;14:20. Published 2015 Jun 6. doi:10.1186/s12942-015-0013-.

